# Function of the auditory cortex characterized by its intrinsic dynamic coactivation patterns estimated in individuals

**DOI:** 10.1101/2025.05.04.651766

**Authors:** M. Hakonen, K. Lankinen, P. Kotlarz, J. R. Polimeni, T. Turpin, J. Ren, D. Wang, H. Liu, J. Ahveninen

## Abstract

Determining the functional organization of the auditory cortex (AC) has been difficult with conventional task-based approaches due to the broad responsiveness of auditory subregions to various acoustic properties. Moreover, most studies have investigated functional organization of AC with static methods, although brain has shown to be organized into dynamic networks. Here, we investigated dynamically varying coactivation patterns of the local networks in the auditory cortex (AC) determined from 7T fMRI data with a novel individualized network-based algorithm. An eight-pattern solution was selected for closer examination based on its high reproducibility of the occurrence rates (r=0.86) and the spatial topography (r=0.79) between sessions. Dynamic AC patterns successfully captured interindividual variability, as indicated by significantly higher variability between than within individuals for the AC pattern occurrence rates and spatial topographies. The coactivation patterns shared similarities between resting-state and auditory-task data, as indicated by the group-level similarity of 0.84 and individual-level similarity of 0.71 in the spatial topographies. Furthermore, the occurrence rates of AC patterns identified in the task data, using pattern templates derived from resting-state data, correlated with specific task contrast regressors. Our results indicate that the AC function can be characterized by a set of dynamically varying coactivation patterns that are consistently observed during resting state and auditory stimulation, and that get synchronized with auditory input. These findings enhance our understanding of the relationship between spontaneous and stimulus-driven activity in the AC and support the development of more time-efficient paradigms for studying its functional organization.

## 1 Introduction

In comparison to the detailed mapping of other sensory domains, the exact functional organization of the human auditory cortex (AC) has been difficult to characterize. Neurons at different parts of AC are driven by a great variety of stimulus attributes (1–3). Exhaustive testing of all potential feature combinations to examine the functional properties of different AC areas is, thus, difficult even in animal models, not to mention human fMRI studies. A typical human neuroimaging study therefore manipulates only a few stimulus dimensions for testing a limited set of hypotheses. This does not allow for controlling the effect of alternative dimensions, which might be even more relevant for the measured response in the given AC area. A small number of recent fMRI studies have approached this problem by modeling responses to large collections of natural sounds (4–6). For example, a 3T fMRI study (5) utilized multivoxel decomposition methods to find a set of canonical components from responses to 165 different natural sounds, corresponding to six different, but partially overlapping, functional subsystems of AC. Analogous methods, which infer the response dimensions from the structure in the data, were also utilized to tease apart AC areas with simple vs. complex spectral preferences from 7fMRI data (4). However, in these approaches the distinct sound objects need to be presented in isolation, using paradigms that require very large numbers of repetitions, over separate imaging sessions, limiting their practical applicability in, for example, characterizing functional abnormalities.

One possible way to address the challenge of defining complex functional brain networks with a limited amount of data is through the analysis of resting-state functional connectivity. At the whole-brain level, this analysis estimates correlations of spontaneous neural activity in the absence of specific tasks or stimuli to investigate the brain’s intrinsic functional organization and its abnormalities under different conditions. Connectivity-based fMRI studies have, for example, shown that brain functional connectomes are highly individual (7) and that they can predict individual behavior and cognition (8–14), as well as the results of clinical interventions (15–17). Recent fMRI studies have also investigated the functional connectivity of AC (18–20). These studies show that AC can be divided into subareas based on its functional connections and that the functional connectivity of AC is individually variable (18–20) and correlates with various auditory dysfunctions (21–23). However, most studies have aimed to quantify functional connectivity between regions as a static measure that is assumed to be constant over time. This may have resulted in an incomplete understanding of AC’s capacity to organize rapidly evolving auditory sequences into coherent percepts.

Studies with fMRI and other modalities (24, 25) have suggested that there is a repertoire of metastable coactivation patterns that are expressed over time (26–28). The fluctuating patterns of interactions between distributed regions appear to be an intrinsic property of mammalian brain organization, which may facilitate the dynamic integration, coordination, and response to internal and external stimuli that are critical for ongoing cognition and behavior (29, 30). A recently developed approach to studying how brain functional connections fluctuate over time is to identify recurring brain coactivation patterns from fMRI data that represent instantaneous network configurations at single time points (31). Analysis of single-volume coactivation patterns has been successfully used to investigate large-scale network dynamics at the whole-brain level (31–35). Interestingly, a recent study (36) also provided evidence that the brain-wide coactivation patterns observed during hand movements can be identified during resting state. If the processing modes of AC could be similarly identified from the resting state data, it could simplify experimental setups which is important, especially in clinical applications.

We adapted a novel Individualized Network-based Single-frame Coactivation Pattern Estimation algorithm (INSCAPE, Fig. 1) (37) to identify metastable “coactivation patterns” in auditory areas of the superior temporal cortices (STC; see, Fig. S1 for specific areas) during rest as well as during auditory and audiovisual stimulation. We estimated the AC patterns from high-resolution 7T fMRI measured in our previous study of 30 healthy volunteers, each of which participated in three 2-hour fMRI sessions (termed hereon “MGH-1mm dataset”, (18)). The generalizability of the AC patterns was tested using the Human Connectome Project 7T fMRI dataset (38). 7T fMRI was used because it offers higher spatial resolution compared to the conventional 3T fMRI without compromising the signal-to-noise ratio (39). Additionally, 7T fMRI more accurately reflects the underlying neuronal activity than 3T fMRI, due to the increased contribution of smaller vessels at the higher field strength (40–42).

**Figure 1.**
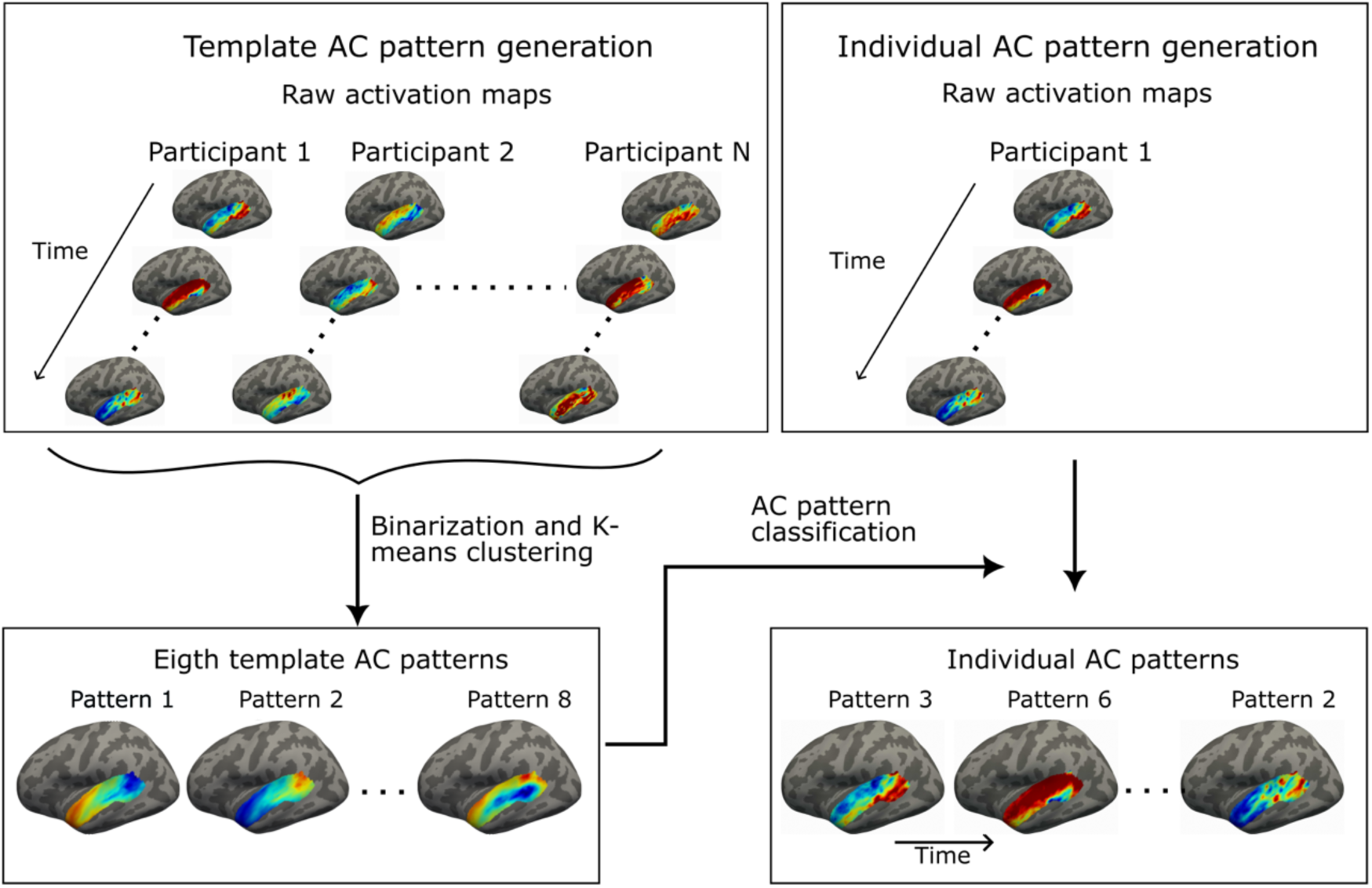
Schematic of the AC pattern generation. The template AC patterns were generated by first binarizing the coactivation maps of each time frame of each participant (i.e., values larger than 0 were set to 1, and values smaller than 0 were set to −1). Thereafter, a k-means clustering algorithm was used to classify the binarized fMRI frames into 8 clusters. Finally, the fMRI frames assigned to each cluster were averaged to create group templates for 8 patterns. The number of patterns was selected based on test-retest reproducibility and visual inspection. At the individual level, each of the fMRI time frames of an individual was classified as the template pattern to which it has the shortest spatial distance.

We hypothesized that coactivation patterns similar to those identified at the whole-brain level could also be reliably identified locally within AC. We further anticipated that the AC patterns identified during the resting state would represent fundamental functional processing modes, showing strong spatial similarity to patterns identified during auditory and audiovisual stimulation. We also expected that the AC patterns would synchronize temporally with specific properties of the auditory and audiovisual input. Finally, we hypothesized the AC patterns to be idiosyncratic, reflected by greater variability in coactivation patterns and occurrence rates between individuals than within individuals.

## 2 Results

### 2.1 Coactivity patterns of AC are reproducible within dataset and generalizable between datasets

The INSCAPE algorithm was used to generate the group template AC patterns (Fig. 1A) using resting-state 7T fMRI data from 30 MGH-1mm participants (two fMRI sessions per participant, 48 min resting-state data in each session). The template patterns were generated by binarizing the coactivation maps of the fMRI timeframes, clustering the binarized maps into clusters across all participants, and averaging the maps within each cluster (Fig. 2A). In the individual-level analysis, each of the fMRI time frames of an individual was classified as the template pattern to which it has the shortest spatial distance (Fig. 1B). The test-retest reproducibility was determined for 2–20 AC pattern solutions by comparing the individual-level mean coactivation maps between the first and second resting-state sessions. The individual-level maps were estimated using group AC pattern templates derived from both resting-state sessions. The eight-pattern solution was chosen for further analysis due to its relatively high test-retest reproducibility at the individual level (0.79, Fig. S1) and the distinctiveness of its spatiotemporal coactivation profiles, as observed through visual inspection. Additionally, the correlation between different coactivation patterns reached a local minimum in the eight-pattern solution (Fig. S2), further indicating that the patterns were sufficiently distinct. However, the test-retest reliability decreased between 4–20 pattern solutions without clear local maxima (Fig. S1). Therefore, other solutions may also be valid. The 10- and 12-pattern coactivation profiles are also shown in the Supplementary Material (Fig. S3).

**Figure 2.**
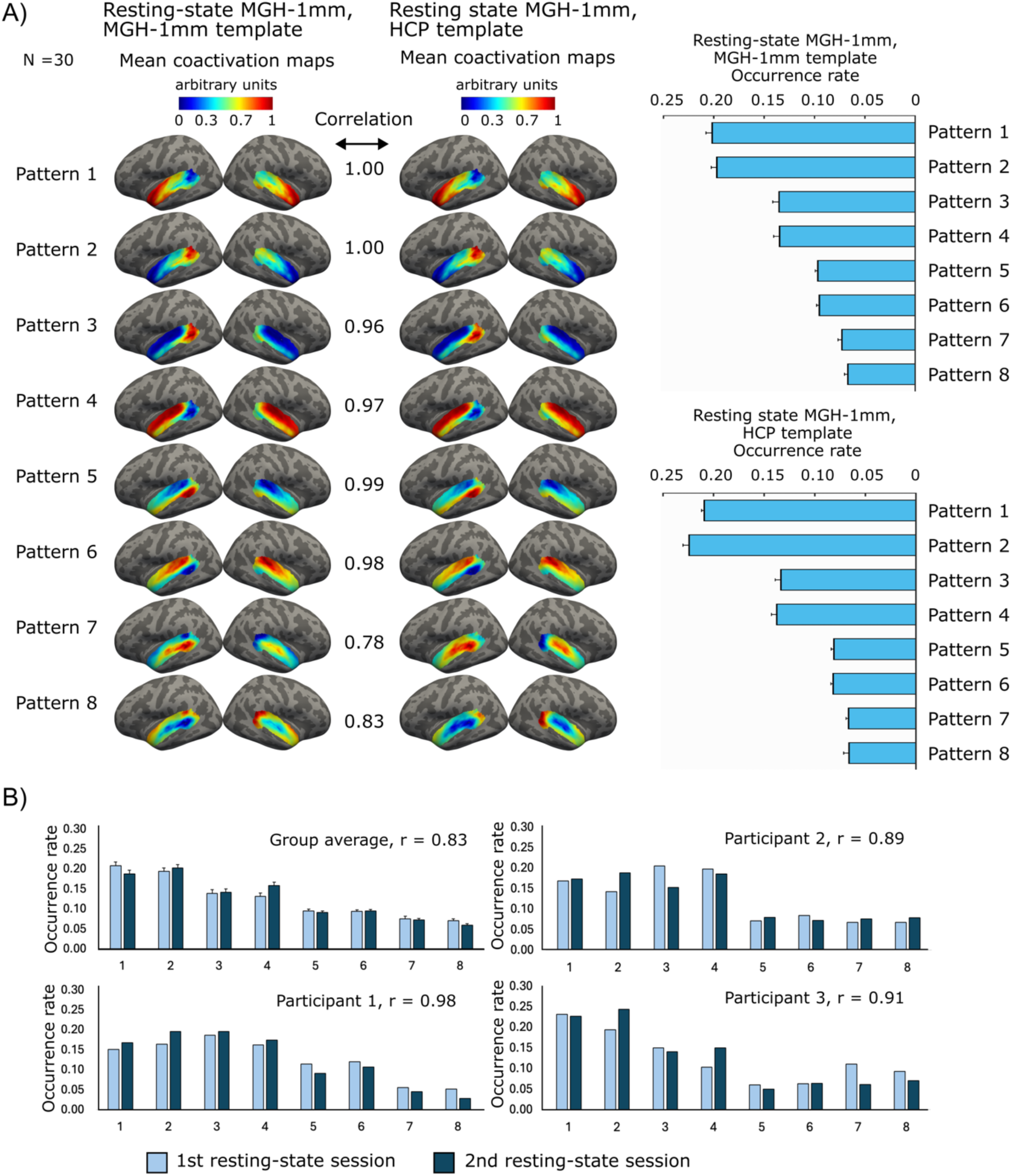
Properties of the AC coactivation patterns. A) The mean coactivation maps and occurrence rates of the AC patterns computed from the resting-state fMRI data of 30 participants measured in our previous study (MGH-1mm data). The maps were calculated using the group templates determined from the MGH-1mm data and the templates determined from the HCP data. The correlation values indicate Spearman correlation between the corresponding group-average patterns of the two datasets. All correlation values were statistically significant (p<0.001 for all patterns). B) AC pattern occurrence rates at the first and second resting-state sessions for group average and three representative participants. Error bars indicate the standard error of the mean (SEM). The group average correlation of 0.83 indicates an average over individual correlations.

The test-retest reproducibility of the AC patterns for the eight-pattern solution was additionally tested by determining both the group template patterns and corresponding coactivation maps and occurrence rates separately from the first and second resting-state fMRI sessions of the MGH-1mm dataset. Each template coactivation pattern was labeled according to the coactivation pattern derived from the whole dataset (Fig. 2A) with which it had the highest spatial correlation. At the group level, the results were highly reproducible, as indicated by a Pearson correlation of 0.96 (p<0.001, Fig. S4) between the mean occurrence rates of the group-level coactivation patterns derived from the two sessions. An average Spearman correlation was 0.96±0.02 (p<0.001 for all patterns) between the group coactivation maps of the corresponding patterns. The significance values were obtained from the Student’s t distribution for Pearson correlation and from the exact permutation distributions for the Spearman correlation (Matlab corr function). The spatial similarity of the activation maps was estimated with Spearman correlation because it estimates if the regions are activated in a similar pattern independently of the exact amplitudes. The results were also relatively reproducible at the individual level, the average Pearson correlation being 0.83±0.04 between the mean occurrence rates. The average Spearman correlation between the coactivation patterns was 0.77±0.02 (patterns: 1: 0.76±0.02, 2: 0.77±0.02, 3: 0.83±0.01, 4: 0.82±0.01, 5: 0.80±0.01, 6: 0.81±0.01, 7: 0.70±0.03, 8: 68±0.03, p<0.001 for all AC patterns). The occurrence rates at the first and second resting-state sessions for the group average and three representative participants are shown in Figure 2B.

To assess the potential impact of spatial dependencies or autocorrelation on the results, we generated coactivation patterns by averaging randomly selected frames from the second resting-state session. Eight patterns were created, each representing an average of frames proportional to the occurrence rate of one of the actual patterns. Subsequently, we calculated the correlation between each actual coactivation pattern derived from the first resting-state session and the corresponding random pattern generated from the second resting-state session. This procedure was repeated 1000 times, and the maximum correlation over the patterns was saved in the null distribution. However, the maximum correlation in the null distribution was 0.41, which is smaller than any of the actual correlations. Thus, the correlations between the actual states cannot be explained with autocorrelation.

Generalizability of the AC patterns was tested by computing the coactivation maps from the MGH-1mm dataset using the group templates derived from the resting-state 7T data from 177 HCP participants (Fig. 2A, right) and comparing them to the coactivation maps obtained with templates derived from MGH-1mm data (Fig. 2A, left). The HCP templates were labeled based on their spatial similarity with the templates derived from the MGH-1mm data. Pearson correlation between the group-level occurrence rates was 0.99 (p<0.001), indicating high generalizability. Spearman correlation between the corresponding group-level coactivation maps derived with these approaches was 0.94±0.09 (p < 0.001 for all patterns, Fig. 2A). We also computed the coactivation maps from the HCP data with templates derived from MGH-1mm and HCP data. Again, the results were highly similar: the correlation between the occurrence rates was 0.99 (p<0.001), and between the corresponding group-level patterns 0.92±0.11 (p<0.001, Fig. S5). Thus, the template patterns determined from different datasets seem to produce consistent results when applied to the same dataset. Additionally, we computed both individual patterns and group templates within MGH-1mm and HCP datasets (Fig. S6). This resulted in a correlation of 0.91 (p<0.002) between the group-level occurrence rates and a correlation of 0.75 between the corresponding coactivation patterns.

Each pattern had an opposite pattern in which the same network was deactivated (Figure 2A: patterns 1 and 2; 3 and 4; 5 and 6; and 7 and 8). Thus, eight patterns represented the activation or deactivation of four networks. There were no differences in the occurrence rates between the patterns and their opposite patterns (p=0.19–1, pairwise Wilcoxon signed rank test, corrected for multiple comparisons), except for the patterns 7 and 8 (p<0.004, pairwise Wilcoxon signed rank test, corrected for multiple comparisons). However, the difference between patterns 7 and 8 was very small: 7.3% of the frames were classified as Pattern 7 and 7.0% as Pattern 8. The occurrence rates were different between the patterns and all other patterns than their opposite patterns (p<0.001, pairwise Wilcoxon signed-rank test, corrected for multiple comparisons).

### 2.2 Occurrence rates and spatial topographies of AC coactivation patterns are more consistent within than between individuals

To investigate the uniqueness of the AC patterns for each individual, the pattern occurrence rates and coactivation patterns were compared between all participants using the resting-state sessions of MGH-1mm data. The similarity between participants was further compared to the similarity between the resting-state sessions within participants to estimate which part of the interindividual variability could be explained by the noise-related variability between sessions. The group templates were determined using both resting-state sessions, and the individual-level AC patterns were derived separately from the sessions.

The mean similarity of the occurrence rates for the eight AC patterns between participants was 0.66±0.32 (i.e. interindividual variability of 0.34). Within participants, the mean similarity was 0.86±0.22, indicating relatively high reproducibility. The occurrence rates were more similar within than between participants (z = 4.8, ranksum = 11064, p < 0.001), indicating meaningful individual variability in the AC pattern occurrence rates (Fig. 3). For the 10-pattern solution, the similarity was 0.81±0.30 within and 0.56±0.35 between participants (z = 5.0, ranksum = 11273, p < 0.001), and for the 12-pattern solution 0.78±0.22 within and 0.42±0.37 between participants (z = 5.9, ranksum = 11903, p < 0.001).

**Figure 3.**
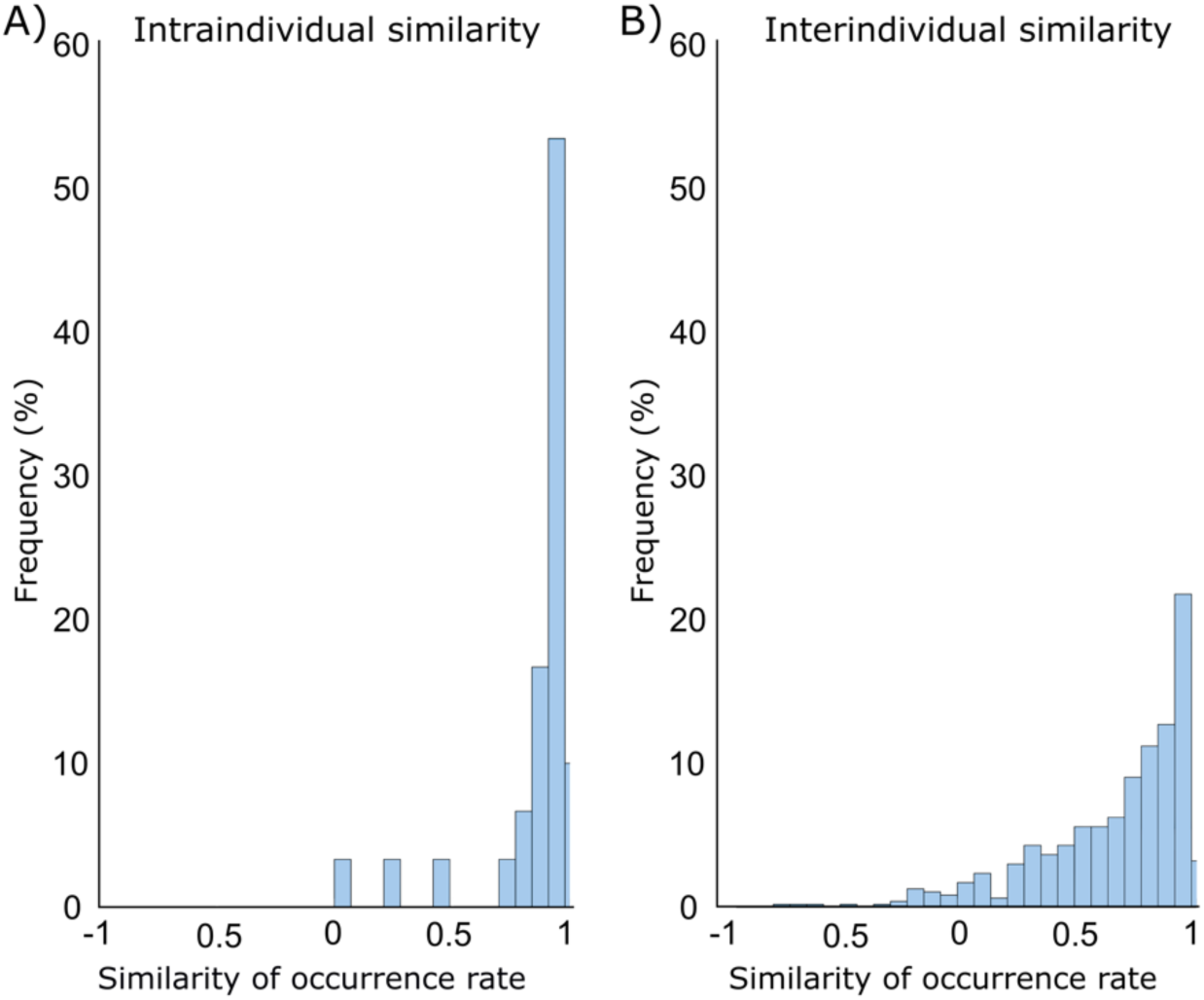
Interindividual variability of AC brain patterns. A) The frequency distribution illustrates the similarity of AC pattern occurrence rates between the first and second resting-state sessions within participants. B) The frequency distribution illustrates the similarity of AC pattern occurrence rates between participants.

The mean similarity of the AC coactivation pattern maps between participants was 0.58±0.08. Within participants, the corresponding similarity was 0.79±0.06, indicating moderate reproducibility. Again, the similarity was higher within than between participants (z = 8.4, ranksum = 13803, p < 0.001). For the 10-pattern solution, the within and between individual similarities were 0.77±0.07 and 0.57±0.08 (z = 8.2, ranksum = 13678, p < 0.001), and for the 12-pattern solution 0.76±0.06 and 0.56±0.08 (z = 8.2, ranksum = 13681, p < 0.001).

The surface-based anatomical normalization methods used in this study (43) have shown to be less accurate in the areas that are not closely related to major sulci and gyri (44, 45). Therefore, individual variability of the spatial topographies of the coactivation maps may have been affected by macroanatomical variability between participants. To examine the potential influence of macroanatomy on our results, we performed a Mantel test to assess whether between-participant similarity in activation topographies correlates with between-participant similarity in AC folding patterns or cortical thickness. The Mantel test indicated no significant correlation in the similarity between coactivation topography and macroanatomy within the AC (curvature: r = −0.04, p = 0.62; thickness: r = −0.14, p = 0.88). This suggests that individual differences in coactivation maps cannot be attributed solely to anatomical variability.

### 2.3 AC coactivation patterns share similarities during resting-state and auditory stimulation

We hypothesized that intrinsic and task-evoked AC coactivation patterns share common network configurations. To test this hypothesis, we determined the template patterns and individual-level coactivation maps from the data measured during auditory and audiovisual tasks and compared the results to the individual coactivation maps derived from the resting-state data using resting-state group templates. The tasks included “Audiovisual Speech/Noise task”, where the participants were presented with clear or blurred video clips of a person voicing either “rain” or “rock.” The auditory component in the videos was either acoustically intact or replaced with noise matched to the spectrotemporal characteristics of the original speech. The other task was “Tonotopy/Amplitude Modulation (AM) task” in which the participants were presented with serial blocks of sounds that varied in their center frequency and amplitude modulation rate.

In Figure 4, the correlation maps derived from the task data are ordered based on their spatial correlation with the maps derived from the resting-state data. The average Spearman correlation between the mean AC coactivation maps derived using these approaches was 0.84±0.20, indicating similarities between the coactivation maps (Fig. 4 A). Especially the first six patterns were highly similar between the approaches (mean correlation 0.94±0.03). The order of the occurrence rates of the pattern pairs 3&4 and 5&6 was opposite between the two approaches. The occurrence rates of pattern pairs 3&4 and 7&8 were closer to each other when the data was measured during auditory stimulation. The differences in the occurrence rates may reflect the engagement of specific networks during the processing of the auditory information.

**Figure 4.**
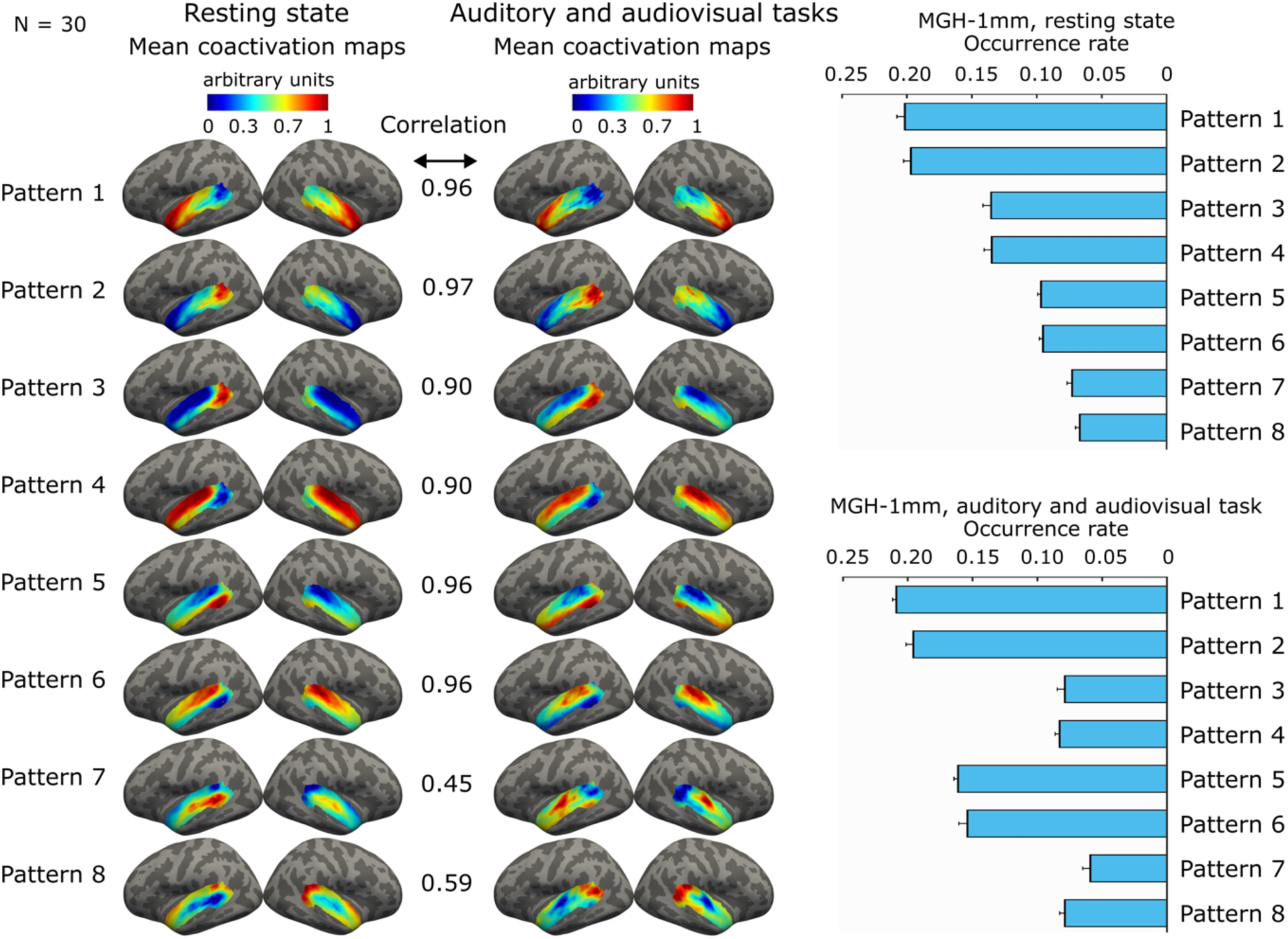
Comparison of occurrence rates and coactivation maps between AC patterns derived from the resting and auditory task fMRI data. The correlation values indicate a correlation between group-average coactivation maps. The resting-state maps were determined using the templates derived from the resting-state data and the task maps using the templates derived from the task data. Error bars indicate the standard error of the mean (SEM).

We also tested whether group templates determined from resting-state data could be generalizable enough to identify similar coactivation maps from the task data that were obtained using templates derived from the task data. The AC coactivation maps derived from the data measured during auditory stimulation are shown in Fig. S7. The mean spatial similarity of the task coactivation patterns obtained using resting-state and task templates was 0.88±0.14. Again, the similarity was highest between the first six patterns (0.96±0.04). The occurrence rates obtained with resting-state vs. task templates from the task data were close to each other.

We also investigated the similarity between the AC patterns derived from the resting-state and task data at the individual level. At the individual level, the correlation of the spatial coactivation maps between task and resting-state data was 0.71±0.07 (p < 0.001, z = 4.8, signed rank = 465; Pattern 1: 0.67±0.02; Pattern 2: 0.67±0.02; Pattern 3: 0.73±0.02; Pattern 4: 0.75±0.02; Pattern 5: 0.73±0.01; Pattern 6: 0.73±0.01; Pattern 7: 0.61±0.02; Pattern 8: 0.60±0.03).

### 2.4 Occurrences of AC coactivation patterns correlate with auditory and audiovisual stimulation

To study if the coactivation patterns derived from the resting-state fMRI could be associated with the AC function, we calculated the correlation between the group-averaged occurrence rates of the AC patterns and 11 auditory and audiovisual task contrast regressors (Fig. 5). For this analysis, the occurrence rates were determined from the task fMRI data using the pattern templates derived from the resting-state data.

**Figure 5.**
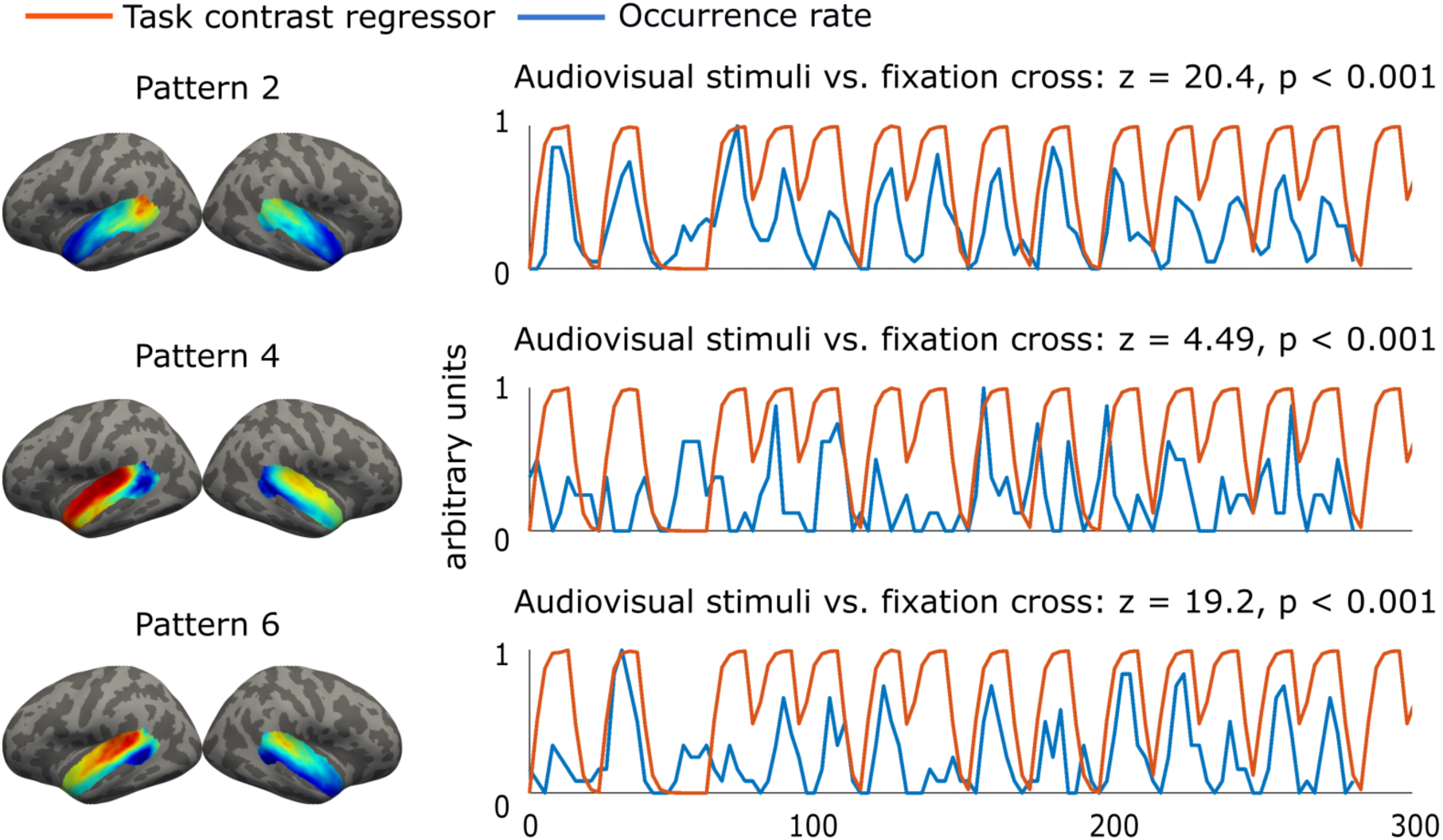
Occurrence rates of the AC coactivation patterns correlate with auditory and audiovisual task regressors. The occurrence rates of three AC coactivation patterns and the task contrast regressors indicating audiovisual stimulation vs. fixation cross.

Each coactivation pattern correlated positively or negatively with 2–9 task contrast regressors (Fig. 5, Table 1). If the pattern correlated positively with a given contrast, its opposite correlated negatively or not significantly with the same contrast. Four AC patterns (2, 4, 6, and 7) showed positive correlations with 4–8 task contrasts. Their opposite patterns (1, 3, 5, and 8) correlated negatively with 2–5 task contrasts and positively with only 1–3 contrasts or not with any contrast.

**Table 1:** z-values of the correlations between the task contrast regressors and coactivation pattern occurrence rates. Aud vs. base = auditory vs. baseline; Hi vs. fast freq. = high (1.87 kHz or 7.47 kHz) vs. low (0.12 kHz or 0.47 kHz) carrier frequencies; Slow vs. fast AM = slow (4 cycles/s) vs. fast (32 cycles/s) amplitude modulation; AV vs. fix = audiovisual stimuli vs. fixation cross; Clear vs. blurred = clear vs. blurred video clips; noise-clear AVia = audiovisual interaction in which the processing of noisy auditory information is enhanced with visual information; AVia = audiovisual interaction in which processing of blurred visual information is enhanced with auditory information; An = noisy auditory stimulus; Vn = noisy visual stimulus; Ac = clear auditory stimulus; Vc = clear visual stimulus.

| Contrasts | AC coactivation patterns |  |  |  |  |  |  |  |
| --- | --- | --- | --- | --- | --- | --- | --- | --- |
|  | 1 | 2 | 3 | 4 | 5 | 6 | 7 | 8 |
| Aud vs. base | -5.97*** | 7.12*** |  | 4.45*** | -10.1*** | 6.99*** |  |  |
| Hi vs. low freq. |  | 2.75* |  |  |  |  |  |  |
| Slow vs. fast AM | -4.51*** | 2.81* |  |  |  | 4.37*** | 3.40** | -3.39** |
| AV vs. fix | -19.0*** | 20.4*** | -3.73** | 4.49*** | -22.6*** | 19.2*** |  | 2.78* |
| Speech vs. noise |  |  | -6.58*** | 2.23* |  | 3.18** | 4.41*** | -8.02*** |
| Clear vs. blurred |  | 2.61* | 3.37** | -3.73** |  |  |  |  |
| noise-clear AVia |  | 3.02** | 2.69* | -2.92** |  | -2.61* |  |  |
| AVia |  | -3.02** | -2.69* | 2.92** |  | 2.61* |  |  |
| AnVcAnVn |  | 3.99*** | 4.30*** | -4.74*** |  | -2.48* |  |  |
| AcVnAnVn |  | 3.07** | -2.70* |  |  |  | 3.77** | -5.87*** |
| AcVcAnVc |  |  | -6.59*** | 3.65** |  | 4.11*** | 2.46* | -5.37*** |

The occurrence rates of Patterns 2, 4, and 6 had the strongest positive correlations with the main effects indicating the processing of audiovisual information vs. baseline and auditory processing vs. baseline (z=4.45–20.4, p<0.001, Table 1). The occurrence rates of patterns 4, 6, and 7 correlated positively with the regressors, indicating the contrast between clear vs. noisy speech (z=2.23–4.41, p<0.05). Moreover, the occurrence rates of patterns 2 and 3 correlated positively with the regressors related to the processing of visual information and audiovisual interaction in which auditory information enhances the processing of blurred visual information (z=2.61–4.30, p<0.05).

## 3 Discussion

This study shows that reoccurring coactivation patterns can be reliably identified within AC from resting-state 7T fMRI data with an individualized network-based algorithm. The coactivation patterns varied meaningfully between individuals both in their occurrence rates and spatial topographies. Interestingly, the coactivation patterns shared similarities between the resting-state data and data measured during auditory and audiovisual tasks. Moreover, the occurrence rates of the AC patterns correlated with specific task contrast regressors, which reflected functions in which the activated network of the pattern had been associated in previous studies, such as processing of auditory features, speech processing, and visual processing. Each coactivation pattern had an opposite pattern in which the same network was deactivated. These results extend previous literature on auditory neuroscience that has mainly focused on static functional connectivity by showing that AC function can be characterized by a dynamically varying individual-specific set of coactivation patterns. Our study also advances studies of dynamic functional connectivity by indicating that coactivation patterns can be identified within local AC networks, and they are, thus, not specific to brain-wide networks.

### 3.1 AC’s intrinsic activity can be characterized by a set of coactivation patterns

An increasing number of fMRI studies have identified dynamically reoccurring brain-wide coactivation patterns demonstrating that functional interactions between brain regions are not static over time but rather highly dynamic. The dynamically varying network configurations have been identified both during resting state and various dynamic tasks (46, 47). Thus, fluctuating patterns of interactions between distributed regions appear to be an intrinsic property of mammalian brain organization, which may facilitate the dynamic integration, coordination, and response to internal and external stimuli that are critical for ongoing cognition and behavior (29, 30). Our results extend these previous studies of dynamic connectivity that have mainly focused on brain-wide networks by showing that dynamic coactivation patterns can also be reliably identified within local networks of AC from single time frames of the resting-state fMRI data. Furthermore, these results expand the auditory neuroscience research that has usually computed average functional connectivity across the whole scan, assuming connectivity to be static over time.

We determined AC coactivation patterns from the single time-frames of 7T resting-state fMRI data using the INSCAPE algorithm (37). The eight-pattern solution was selected for closer examination due to its relatively high individual-level test-retest reproducibility (0.79). The reliability declined across the 4–20 pattern solutions range without clear local maxima (Fig. S1), indicating that alternative solutions may also be valid. The spatiotemporal coactivation profiles of the eight pattern-solution also differed based on the between-pattern spatial correlations (Fig. S2). The individual-level reproducibility between the occurrence rates was 0.86 which is lower than 0.90 reported previously for the whole brain (37). Our results demonstrate that meaningful coactivation patterns can be identified within AC even without any a priori information provided by weighting with a group atlas, as was used in (37). Furthermore, the reproducibility can be expected to be lower for small AC networks than large-scale networks of the whole brain that may have a lower signal-to-noise ratio. At the group level, the patterns were highly reproducible, as demonstrated by the average correlation of 0.96±0.02 between the coactivation patterns and one between the cooccurrence rates of the patterns determined from two fMRI sessions of the same participants.

The AC coactivation patterns were also generalizable, as indicated by highly similar coactivation maps and co-occurrence rates obtained for the dataset with the templates derived from the same or different dataset (Fig. 2, Fig. S5). There was still considerable similarity in the activation maps and their occurrence rates even when both the coactivation patterns and templates were determined independently within datasets (Fig. S6). Related results have been obtained in previous studies that have shown brain-wide coactivation patterns to share spatial similarities between datasets (48–50).

While brain-wide coactivation patterns have usually been shown to be hemispherically symmetric (48, 50–52), the AC patterns showed notable hemispheric asymmetry. This likely reflects the lateralization of the AC functions (53). In contrast, the brain-wide coactivation patterns are usually dominated by the default mode network, known to be hemispherically symmetric. However, with a large number of patterns, hemispheric lateralization has also been identified in brain-wide patterns. For example, a recent study (37) found one of the sixteen patterns to be left-lateralized and one right-lateralized. The occurrence rate of the left-lateralized pattern was also higher during a language task than resting state and correlated with language-task onsets.

### 3.2 Coactivated patterns of AC are organized in pairs with opposite network topographies

The AC patterns formed pairs with opposite coactivation maps. There were no significant differences in the occurrence rates between the patterns and their opposite counterparts. If the pattern correlated positively with a task contrast regressor, its opposite pattern never correlated with the same task contrast or correlated with it negatively.

Opposite coactivation maps have previously been reported for brain-wide networks (48, 50–52). Some opposite activation maps were also found in the previous study that identified sixteen whole-brain patterns using the INSCAPE algorithm (37). The opposite coactivation patterns may reflect anticorrelated networks commonly found in resting-state functional connectivity studies (54–59). For example, the activity of the default mode network is decreased during the execution of cognitive tasks and increased during the resting state. Thus, the default mode network is also referred to as a task-negative network that often correlates negatively with task-positive networks (55, 60). The strength of the negative correlation between the default mode network and the task-positive network has been associated with interindividual differences in task performance and task-evoked fMRI responses (61). Our results extend these previous studies by demonstrating that anticorrelated networks are not specific to brain-wide networks, but can also be found within local networks of AC.

The meaning of the anticorrelated networks is unclear, but they may represent the separation of neuronal processes linked to competing cognitive demands, such as focusing on a task versus engaging in stimulus-independent thoughts (55). In this study, the anticorrelated AC networks may compete with each other while processing different features of auditory or audiovisual information. A rat study also found opposite patterns with opposite phases, suggesting that they may reflect peaks and troughs of periodic neural oscillations (62). While it has been proposed that these anticorrelations could be artifacts introduced by global signal (GS) regression (63, 64), growing evidence demonstrates the presence of anticorrelation patterns independently of preprocessing steps (55–57, 59). Our study further supports the neurophysiological origin of the anticorrelations since GS was not regressed out from the data.

### 3.3 AC coactivation patterns capture meaningful interindividual variability

Both the occurrence rates (interindividual similarity = 0.66) and spatial topographies (interindividual similarity = 0.58) of the coactivation patterns varied substantially between individuals. Within individuals, the occurrence rates (interindividual similarity = 0.86) and spatial topographies (interindividual similarity = 0.79) of the coactivation maps were more similar. These results indicate that the coactivation patterns are relatively reproducible within individuals but also capture meaningful interindividual variability. It should be noticed that the individual variability of the spatial topographies may also be affected by macroanatomical variability between participants because the normalization methods are less accurate in the areas that are not strongly related to major sulci and gyri (44, 45). However, the between-participant similarity in the spatial topographies did not correlate with the between-participant similarity in macroanatomy within AC (curvature: r = −0.04, p = 0.62, thickness: r = −0.14, p=0.88), suggesting that the individual variability in the coactivation maps is not explained by anatomical variability alone. Our results are in line with two recent studies that showed the brain-wide coactivation maps to vary substantially between individuals (34, 37). Together with these results, our study expands previous studies that have mostly ignored individual variability of the dynamic coactivation patterns. Our results are also in accordance with recent static functional connectivity fMRI studies indicating that the functional organization of AC varies substantially between individuals (18, 20) and that individual variability of the AC functional connectivity is even higher than in the visual cortex (19). Our results show that individual variability can not only be found in static functional connectivity of AC, but also in the occurrence rates and spatial topographies of the dynamically varying network configurations.

### 3.4 Processing of auditory information can be characterized with AC intrinsic coactivation patterns determined from resting-state fMRI data

The AC pattern coactivation maps shared similar network configurations when determined using group templates derived from resting-state and task data. The average correlation of the coactivation maps between the resting-state and task data was 0.84 at the group level and 0.70 at the individual level. These results are in line with the studies that have found only small differences in the static functional connectivity networks between the resting state and task (65–68). One possibility could be that the intrinsic AC patterns determined using the resting-state data represent basic processing units synchronizing with external stimuli. We tested this hypothesis by computing the correlation between the AC pattern occurrence rates and task contrast regressors. The occurrence rates were computed from the task data using the group templates derived from the resting-state data.

The occurrence rates of the AC coactivation patterns correlated with several task contrast regressors, suggesting that they reflect the processing of auditory and audiovisual information within AC. Six of the AC patterns correlated positively with 1–8 regressors, and two AC patterns only negatively. The regressors with which the patterns correlated positively also reflected functions in which the areas activated in the spatial topographies of the patterns were associated in previous studies. The topography of Pattern 2 displayed activation in the area comprising the temporoparietal junction (TPJ) and most posterior parts of the superior temporal gyrus (pSTG), and the activity was stronger in the left hemisphere. The posterior superior temporal cortex (pSTC) has been shown to be highly responsive to sound onsets, including onsets of speech and nonspeech sounds (69). In line with these results, Pattern 2 correlated most strongly with the regressors reflecting auditory stimulation vs. baseline and audiovisual stimulation vs. fixation cross. pSTG has also been shown to be important for audiovisual integration (70, 71), which is again in line with our finding that Pattern 2 correlated with the regressor indicating an interaction where visual input enhances the processing of noisy speech and the regressor indicating clear vs. blurred visual information. Pattern 2 also correlated with clear vs. noisy speech sounds, which agrees with the studies showing TPJ to be activated by language and semantics (72).

Patterns 4 and 6 were correlated with the regressors which reflect auditory information processing, but not with regressors related to visual or audiovisual processing. Patterns 4 and 6 were also correlated with the regressor, indicating Speech vs. noise contrast. The spatial topography of Pattern 4 showed activity in the primary auditory areas that are known to be highly responsive to acoustic information (e.g. frequencies, amplitude modulation) (73–75). In the spatiotemporal topography of Pattern 6, the activity extended from the primary auditory areas to the anterior STC and temporal pole. These areas have been associated with the processing of speech, words, and semantics (73–76). The activity was left lateralized for both Pattern 4 and Pattern 6.

The spatial topography of Pattern 7 showed bilateral activity expanding from the posterior to middle STG. The activity is stronger in the left hemisphere and includes also parts of the posterior superior temporal sulcus (pSTS). Pattern 7 had the strongest correlations for regressors, indicating speech vs. noise and slow vs. fast AM, suggesting its role in speech processing. These results are in accord with previous literature showing pSTG, mSTG, and STS to be activated by words (77–81).

The spatial topography of Pattern 3 displayed left-hemispheric activation in the pSTS. Pattern 3 was correlated with the regressors indicating clear vs. blurred visual processing and an audiovisual interaction where visual information enhances the processing of noisy speech. Pattern 3 did not correlate with any regressors specific to auditory or speech processing. Together, these results suggest that Pattern 3 could be important in the processing of audiovisual and visual information. This conclusion is consistent with studies that have associated left pSTS with the integration of audiovisual information (82–85). The pSTS has also been shown to be activated during lip-reading (86) and dynamic facial expressions (87), which is further in line with our results as the audiovisual stimulus included video clips of a woman voicing “rain” or “rock”.

If the pattern correlated positively with the regressor, its opposite pattern correlated negatively with the same regressor or did not correlate significantly with it. Of the three regressors with which the patterns had the strongest correlations, 2–3 were correlated negatively with the corresponding opposite patterns. This was true for all patterns, expect for Pattern 8 which only had a weak (r=0.09) positive correlation with one regressor. These results are, again, in line with the studies that found anticorrelation within large-scale functional networks (54–59) and further support the conclusion that anticorrelated networks may also exist within local AC networks as discussed in 3.2.

### 3.5 Limitations

While this study provides valuable insights into the dynamically varying networks within the AC, it is important to recognize its limitations. The number of AC patterns was selected based on test-retest reliability. Using additional metrics, such as the Silhouette coefficient, could help to study if there are optimal pattern numbers. Second, while in this pioneering study focusing on AC, we needed to follow previous procedures, a possible approach for further studies would be to collect more data from each participant (34). In this case, the individual variability could be taken into account by determining the template patterns individually which may increase in individual-level reproducibility. Finally, the influence of inadequate registration of cortical surfaces on the individual variability of the AC patterns could be further decreased by using more sophisticated registration methods (44, 45).

### 3.6 Conclusions and future directions

This study had four major findings: 1) AC function can be characterized by a set of dynamically varying coactivation patterns derived from single time-frames of the resting-state 7T fMRI data; 2) AC patterns derived from the resting-state and task fMRI data share a lot of similarities; 3) AC patterns get temporally synchronized with auditory stimulation, and 4) AC patterns capture meaningful interindividual variability. Identification of reproducible and relatively generalizable dynamic coactivation patterns advances previous research of AC functional organization that has mainly focused on static functional connectivity. By indicating that dynamically varying connectivity patterns are also highly individual, our results support and extend studies that have identified high individual variability in static functional connectivity of AC. Additionally, our results show that the dynamic AC network configurations remain relatively unchanged during the resting state compared to the task and get synchronized with specific features of auditory or audiovisual input. An important finding of our study is also that coactivation patterns are not specific to brain-wide networks but can also be found locally within AC. Interesting topics for further studies are associating the AC patterns with a more comprehensive set of auditory stimuli and investigating how individual variability in the AC patterns is reflected in auditory function, hearing abilities, and speech comprehension assessed with behavioral and clinical tests. An important question is also whether auditory processing problems could be identified from the AC coactivation patterns derived simply from the resting state instead of using several different auditory stimulus paradigms.

## 4 Methods

### 4.1 Data

#### 4.1.1 MGH-1mm dataset

The MGH-1mm dataset consisted of the same 7T fMRI data from 30 volunteers (32.4 ± 10 years, 15 women, all right-handed) that was published in our previous study (18). The study protocol was approved by the Institutional Review Board at Mass General Brigham, and all participants provided informed consent prior to the experiments. Twenty-eight of the participants were native English speakers, and none of them reported hearing impairments, exposure to excessive noise, or the use of cognition-altering medications. All except one of 20 participants who participated in the Hughson-Westlake pure tone procedure had hearing thresholds below 25 dB at 0.125–8 kHz (for one participant, 40-dB threshold at 6 and 8 kHz in the left ear and 35 dB at 8 kHz in the right ear).

Twenty-two participants completed three fMRI sessions on different days, while eight completed them in four days. In two sessions, six 8-minute resting-state fMRI scans were conducted (except for one participant, for whom only four scans were performed in the other resting-state session). The tasks were carried out over one or two separate sessions. They included a combined tonotopy and amplitude modulation (AM) rate representation task (2 × 8 min) and an audiovisual speech perception task (4 × 11 min).

MRI data were acquired using a 7T whole-body MRI scanner (MAGNETOM Terra, Siemens, Erlangen, Germany) with a custom-built 64-channel receive brain array coil and a single-channel birdcage transmit coil (88). Each imaging session included: **1)** T1-weighted anatomical images collected using a 0.75-mm isotropic Multi-Echo MPRAGE pulse sequence (89, 90) (TR = 2530 ms; TE = 4 echoes with TEs 1.72, 3.53, 5.34 and 7.15 ms; flip angle = 7°; field of view, FoV = 240 × 240 mm^2^; 224 sagittal slices), **2)** blood-oxygenation-level-dependent (BOLD) fMRI data collected using a single-shot 2D simultaneous multi-slice echo planar imaging (EPI) sequence (91) (blipped-CAIPI, acceleration factor in slice-encoding direction: 3; acceleration factor in phase-encoding direction: 4; TR = 2800 ms; TE = 27.0 ms; 1-mm^3^ isotropic voxels; flip angle = 78°; FoV = 192 × 192 mm^2^; 132 axial slices; phase enc. dir.: anterior→posterior; readout bandwidth = 1446 Hz/pixel; nominal echo spacing = 0.82 ms; fat suppression), **3)** a short EPI scan (three repetitions) collected with the same parameters, but with the opposite phase-encoding direction polarity (posterior→anterior, PA-EPI), and **4)** a gradient-echo based B_0_ field map (TR = 1040 ms, TEs = 4.71 ms and 5.73 ms; 1.3-mm^3^ isotropic voxels; flip angle: 75°; FoV: 240 × 240 mm2; 120 slices; bandwidth = 303 Hz/pixel). In addition, T2-weighted anatomical images were acquired for 28 out of 30 participants with the T2 SPACE sequence (voxel size = 0.83 × 0.83 × 0.80 mm, TR = 9000 ms, TE = 269 ms, flip angle = 120°, FoV = 225 × 225 mm^2^, 270 sagittal slices) in one session. The different contrast of T2 images was used to improve pial surface reconstruction. During the fMRI data acquisition, participants’ heart rate and respiration signals were recorded at 400 samples/s using the Siemens photoplethysmogram transducer and respiratory belt.

#### 4.1.2 Human Connectome project 7T fMRI dataset

The generalizability of the AC coactivation patterns was tested using resting-state 7T fMRI and structural 3T MRI data measured from 184 participants (age range: 22–35, the ages were reported in age bands of 3-4 years, two participants were over 36 years old, 112 women) in the Human Connectome Project (HCP, https://www.humanconnectome.org/hcp-protocols-ya-7t-imaging). The structural data was acquired using a customized Siemens 3T Connectome Skyra scanner with a standard 32-channel Siemens receive head coil. T1-weighted anatomical images were collected using a MPRAGE pulse sequence (accel. factor PE = 2; multi-band accel. Factor = 5; TR = 2400 ms; TE = 2.14 ms; 0.7 mm^2^ isotropic voxels; flip angle = 8°; field of view, FoV = 224× 224 mm^2^; 256 slices). The resting-state data were acquired using a Siemens Magnetom 7T MR scanner with the Nova32 32-channel Siemens receive head coil and an incorporated head-only transmit coil. The total amount of the resting-state data measured in each session was approximately 16 min. The participants were instructed to keep their eyes open with a relaxed fixation on a projected cross on a dark background. Oblique axial slices were acquired in a posterior-to-anterior direction in runs 1 and 3 and in an anterior-to-posterior phase encoding direction in runs 2 and 4. The fMRI data was acquired with the gradient-echo EPI sequence (accel. factor PE = 2; multi-band accel. factor = 5, TR = 1000 ms, TE = 22.2 ms, 1.6 mm^3^ isotropic voxels, flip angle = 45°, FOV = 208 x 208 mm^2^, 85 slices, readout bandwidth = 1924 Hz/pixel; nominal echo spacing = 0.64 ms).

### 4.2 Preprocessing

The MGH-1mm dataset was preprocessed in our previous study, which also describes the preprocessing pipeline in more detail (18). Preprocessing of the structural images consisted of the bias field correction and automatic generation of the cortical surfaces using the recon-all function of FreeSurfer 7.1 (92) with an extension for submillimeter 7T data (89). Both the T2 image and an average of 3–4 T1 images from different sessions were used in the reconstruction to improve the quality of the cortical surfaces. The preprocessing of the fMRI data included slice time correction, rigid body correction of head movements, geometric distortion correction, correction of heart rate and respiratory artifacts with RETROspective Image CORrection (RETROICOR) algorithm (93), boundary-based registration of the functional data with the anatomical images, intracortical smoothing (94), band-pass filtering between 0.01–0.08 Hz, and nuisance signal regression of head-motion parameters, ventricular and white-matter signals, and signals outside of the head.

We downloaded the HCP structural MRI data that had already been processed with the PreFreeSurfer pipeline (95). This included gradient nonlinearity distortion correction, coregistration and averaging of the T1 run and T2 run, alignment along the anterior and posterior commissure (AC–PC) axis, brain extraction, field map distortion correction, registration of T2 to T1, bias field correction, and MNI nonlinear volume registration. We generated the cortical reconstructions from these data for each participant using the recon-all function of FreeSurfer 7.1 (92). The T2 anatomical data were used to improve the outcome for pial surface reconstruction in the recon all algorithm. We used the fMRI data that had already been processed with an HCP minimal preprocessing fMRI volume pipeline described in detail in (95). The minimal preprocessing included motion correction using 6-DOF FLIRT, gradient nonlinearity distortion correction, and EPI to T1w anatomical registration. The original EPI frames were resampled to the atlas space via one-step spline resampling that included all transforms (motion correction, EPI distortion correction, EPI to T1w with FLIRT BBR, fine-tuning of EPI to T1w with bbregister, nonlinear T1w to MNI). The bias field estimated from the structural data was also removed from the fMRI data. The fMRI data was masked and normalized to a 4D whole-brain mean intensity of 10,000. We resampled these preprocessed volumes into fsaverage6 surface and regressed out head-motion parameters and ventricular and white-matter signals. The data were also spatially smoothed with a 3D 6-mm full-width at half-maximum Gaussian kernel.

### 4.3 Single-frame dynamic coactivation analysis

The INSCAPE analysis starts with the generation of the group-level template coactivation patterns that are thereafter used to estimate individual-specific patterns (37). To create the template patterns, fMRI frames were first binarized so that positive values were set to 1 and negative values to −1. The binarized fMRI frames of all participants were clustered into eight clusters using the k-means algorithm. Finally, the binarized frames were averaged across participants in each cluster. The resulting eight averaged maps were used as group templates of AC patterns. Eight patterns were selected because their spatial topographies had relatively high test-retest reliability (0.79), and their spatial maps were different from each other (Fig. S2). However, there were no clear local maxima in the reliability (Fig. S1), and other solutions may also be valid. To minimize the effect of motion artifacts on the templates, the templates were determined only using the participants with a mean and maximum head motion (framewise displacement) under 0.1 and 0.5 mm, respectively. Of the total 60 sessions (two sessions for each of the 30 participants), 55 fulfilled this criterion.

We also attempted to weigh the raw activation maps with parcellation confidence derived from group AC parcellations consisting of 4 and 6 functional networks (18), correspondingly as was done for the whole-cortex analysis in the previous study (37). However, the resulting spatial topography maps contained no clear, continuous, larger activated/deactivated networks. This may be because the networks in the parcellations were much smaller than the ones used for the whole brain (37). The group-level networks may not have aligned with the individual-level networks because of the interindividual variability. Therefore, we decided not to weigh the raw activation maps with the parcellation confidence in contrast to the INSCAPE procedure presented in the previous study (37).

The individual-specific AC patterns were estimated by comparing each fMRI time frame to the template patterns and assigning it to the pattern with the shortest spatial distance (cosine similarity, MATLAB, pdist function). The time frames classified as the same AC pattern were averaged to create an individual-level AC pattern map. At the individual level, the frames were not binarized. The pattern occurrence rates were determined as the number of time frames assigned into the same AC pattern divided by the total number of time frames. Each of the coactivation patterns was bilaterally normalized between 0–1 for visualization.

### 4.4 Reproducibility and generalizability of AC patterns

The reproducibility of the AC patterns was estimated by deriving the template and individual AC patterns separately from the first and second resting-state sessions of the MGH-1mm dataset. Both group average and individual level coactivation map topographies were compared between the two sessions with Spearman correlation and the occurrence rates with Pearson correlation. At the individual level, the AC pattern similarity within and between participants was compared using the Wilcoxon rank sum test. The correlation values were transformed into z-scores with Fisher’s z transformation for the Wilcoxon rank sum test.

To estimate the generalizability of the AC patterns, the template AC patterns were derived separately from the resting-state MGH-1mm data and HCP data. Thereafter, the individual-level AC patterns were determined from the MGH-1mm data using the MGH-1mm and HCP templates. Finally, the group-average AC pattern topographies and occurrence rates were derived using MGH-1mm compared to the ones derived using HCP templates. This allowed us to see whether the AC templates derived from different datasets produce similar results when applied to the same data. The same analysis was conducted by deriving the individual-level patterns from the HCP data. Additionally, we tested the generalizability of the AC patterns by deriving both the AC template patterns and individual-level patterns within MGH-1mm and HCP datasets and comparing the group-average results.

To evaluate the potential influence of spatial dependencies or autocorrelation on our results, we generated eight coactivation patterns by averaging randomly selected frames from the second resting-state session of the MGH-1mm dataset. Each random pattern was created by averaging a number of frames proportional to the occurrence rate of its corresponding actual pattern. We then calculated the correlation between each actual coactivation pattern derived from the first resting state session and its corresponding random pattern from the second session. This process was repeated 1,000 times, and the maximum correlation across patterns was used to construct the null distribution. Finally, the correlations between the actual patterns derived from the first and second resting state sessions were compared against the null distribution.

### 4.5 Interindividual variability of AC patterns

The interindividual variability in the coactivation patterns and their occurrence rates were estimated with pairwise comparisons between participants using resting-state MGH-1mm data. The similarity of the spatial topographies was estimated using the Spearman correlation and the similarity of the occurrence rates with the Pearson correlation. The correlation values were transformed into z-scores with Fisher’s z-transformation. Thereafter, the similarity between participants was compared to the similarity between two resting-state sessions within participants using the Wilcoxon rank sum test. Comparison to the within-participant similarity allowed us to estimate what part of the interindividual variability could be explained with noise.

The Mantel test (96) was further used to test whether interindividual variability in the pattern spatial topographies reflects cortical thickness or curvature variability. The thickness and curvature maps for each participant were created using the FreeSurfer recon-all pipeline, and the AC area used in the AC patterns was extracted from the thickness and curvature maps (see (18) for more details). Similarity matrices for cortical curvature and thickness were created by calculating pairwise Spearman correlations between participants using vectors representing their AC thickness and curvature, respectively. A similarity matrix for the AC pattern topographies was created using pairwise Spearman correlations between participants using vectors representing the pattern topographies. The similarity matrix was computed by averaging the pattern-specific similarity matrices. In the Mantel test, the Spearman correlation was computed between the upper triangle elements of the AC pattern topography similarity and thickness similarity matrices, as well as between AC pattern topography and curvature similarity matrices. The statistical significance was assessed by recalculating the correlations using 10,000 random permutations of the rows and columns in the AC pattern topography similarity matrix relative to each other. The results were corrected for multiple comparisons using the Benjamini-Hochberg procedure with a significance threshold of 0.05 (97, 98).

### 4.6 Correlation between the occurrences of AC coactivation patterns and task contrast regressors

To test whether AC coactivation patterns could be associated with AC function, we calculated correlations between the AC pattern occurrences and task contrast regressors. The vectors indicating AC pattern occurrences were created by classifying each time frame of task fMRI data of each individual into one of the eight AC patterns. The time frames were classified by assigning them to the group template pattern with the shortest spatial distance to it (cosine similarity, MATLAB, pdist function). The group template patterns were derived from the resting-state data. The analysis was conducted using the group average over the individual-specific occurrence rate vectors.

We generated regressors for eleven task contrasts. For the Audiovisual Speech / Noise task, the contrast regressors were computed across the Auditory Clear / Visual Clear (AcVc), Auditory Clear / Auditory Noisy (AcVn), Auditory Noisy / Visual Clear (AnVc), and Auditory Noisy / Visual Noisy (AnVn) stimuli. In the main effect of the Audiovisual vs. fixation cross, we contrasted all possible audiovisual combinations with the fixation cross. In the main effect of Speech vs. Noise, all possible audiovisual combinations were contrasted with clear auditory signal with those with noisy auditory signal, i.e., ((AcVc+AcVn) − (AnVc+AnVn)) / 2. In the main effect of Lip motion vs. Noise, all audiovisual combinations with clear visual signal were contrasted with those with blurred vision, i.e., ((AcVc+AnVc) − (AcVn+AnVn)) / 2. We also computed an audiovisual interaction to identify the cortical areas where visual information of lip motion has the strongest effect on processing speech sounds when the auditory signal is noisy. This contrast was, thus, defined as (AnVc-AnVn) − (AcVc-AcVn). Additionally, we computed an audiovisual interaction for identifying cortical areas where auditory information has the strongest influence on the processing of visual information when the video clip is blurred. This contrast was defined as (AcVc-AcVn) − (AnVc-AnVn). Furthermore, we generated regressors indicating the following contrasts: AnVcAnVn, AcVnAnVn, and AcVcAnVc. From the Tonotopy/AM task, the following contrasts were selected: 1) auditory stimulation vs. baseline, 2) high (1.87 kHz or 7.47 kHz) vs. low (0.12 kHz or 0.47 kHz) carrier frequencies, and 3) slow (4 cycles/s) vs. fast (32 cycles/s) amplitude modulation.

Pearson correlation was computed between the occurrence rate vectors and each of the task contrast regressors using MATLAB R2023a. Thereafter, the correlation values were transformed into z-values by using Fisher’s transformation and further to z-scores by dividing the z-values with the standard error of the z-values. The significance values for the z-scores were determined from the z-distribution and corrected for multiple comparisons with the Benjamini-Hochberg procedure (97, 98). Only results exceeding the significance threshold of 0.05 were reported.

## Acknowledgments

Our work was funded by NIH grants K99DC022305, R011DC016915, R01DC016765, R01DC017991, S10OD023637, R01DC016765, R01DC017915, R01DC017991, P41EB015896 and in part by the NIH grant P41-EB030006. In addition, the work was supported by the MGH/HST Athinoula A. Martinos Center for Biomedical Imaging and was made possible by the resources provided by NIH Shared Instrumentation Grants S10-OD023637. The content is solely the responsibility of the authors and does not necessarily represent the official views of the National Institutes of Health. The project was also funded by Canadian Institutes of Health Research (CIHR MFE-171291), Changping Laboratory (2021B-01-01) and China Postdoctoral Science Foundation (2022M720529). We would like to thank Azma Mareyam for hardware support and Dr. Danny Park for pulse sequence support. Much of the computation resources required for this research was performed on computational hardware generously provided by the Massachusetts Life Sciences Center (https://www.masslifesciences.com/).

## Notes

### Competing Interest Statement

The authors have declared no competing interest.

