## Supplementary material for "Function of the auditory cortex characterized by its intrinsic dynamic coactivation patterns estimated in individuals"

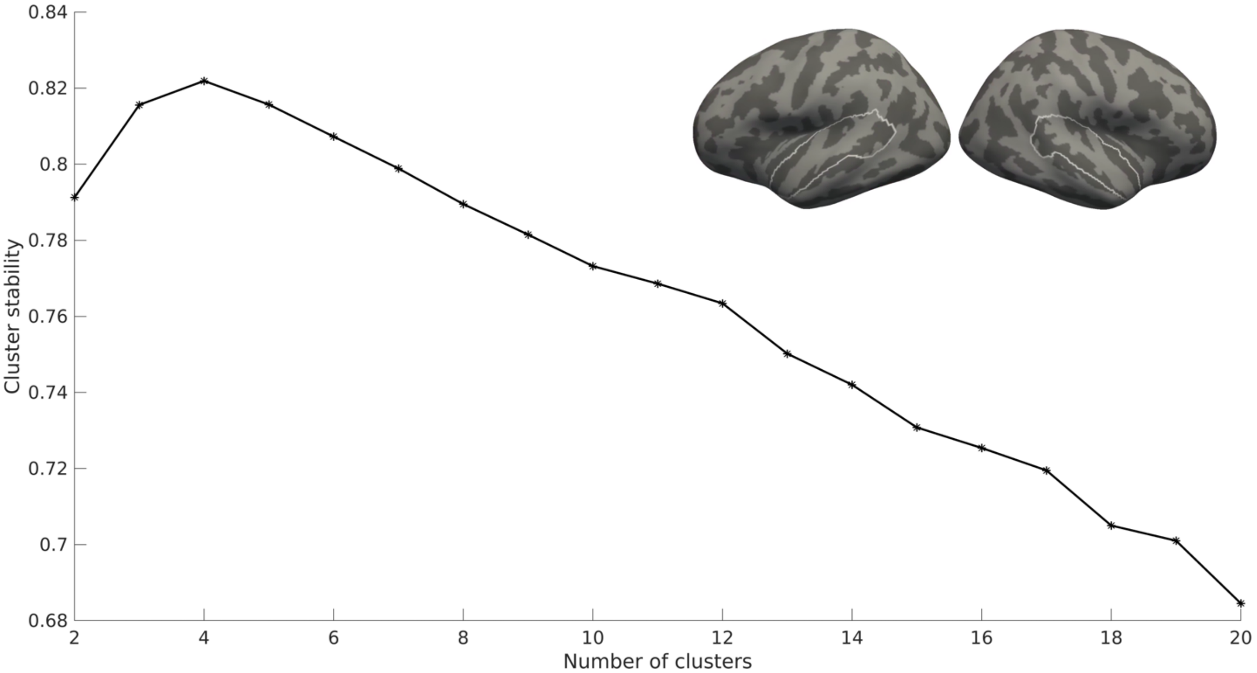


**Figure S1**: Test-retest stability of brain patterns as a function of the number of clusters in the MGH-1mm dataset. Within-participant test-retest stability was estimated as an average spatial similarity of brain patterns between the first and second resting-state sessions. The stability of the cluster analysis was estimated as an average spatial similarity between the brain patterns derived from the first and second resting-state sessions. The group template patterns were determined using the data from both sessions. An 8-cluster solution was selected since it had relatively high stability (0.79), and the spatial profiles of the AC pattern coactivation maps were different from each other based on visual inspection. The AC areas selected for the analysis are outlined on the fsaverage surfaces.

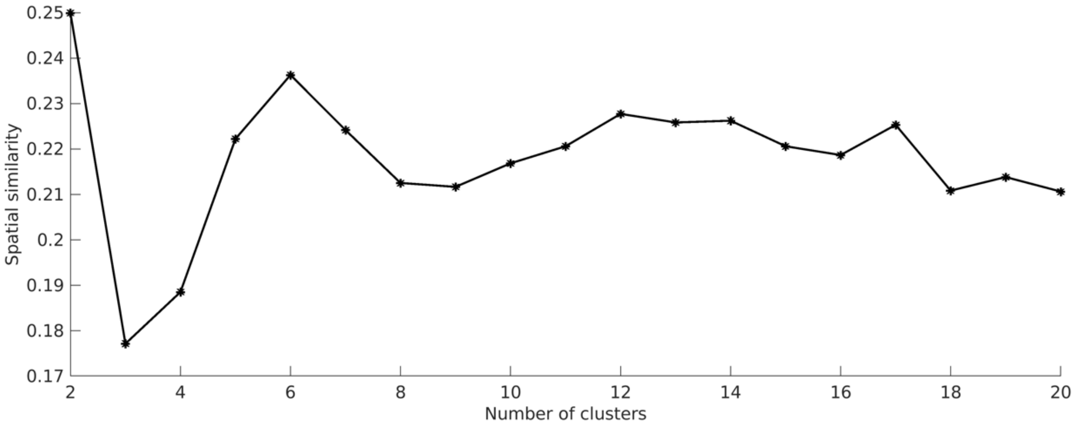


**Figure S2**: Spatial correlation between different AC coactivation patterns as a function of the number of coactivation patterns. The similarity between the AC coactivation patterns was estimated for each solution by averaging the Spearman correlations between all coactivation patterns. The 8-coactivation pattern solution was selected for further analyses since the between-pattern similarity started to increase after that. Since the patterns were represented in pairs with inverse topographies (i.e., the same network was either activated or deactivated), we selected an even number of states for further analysis.

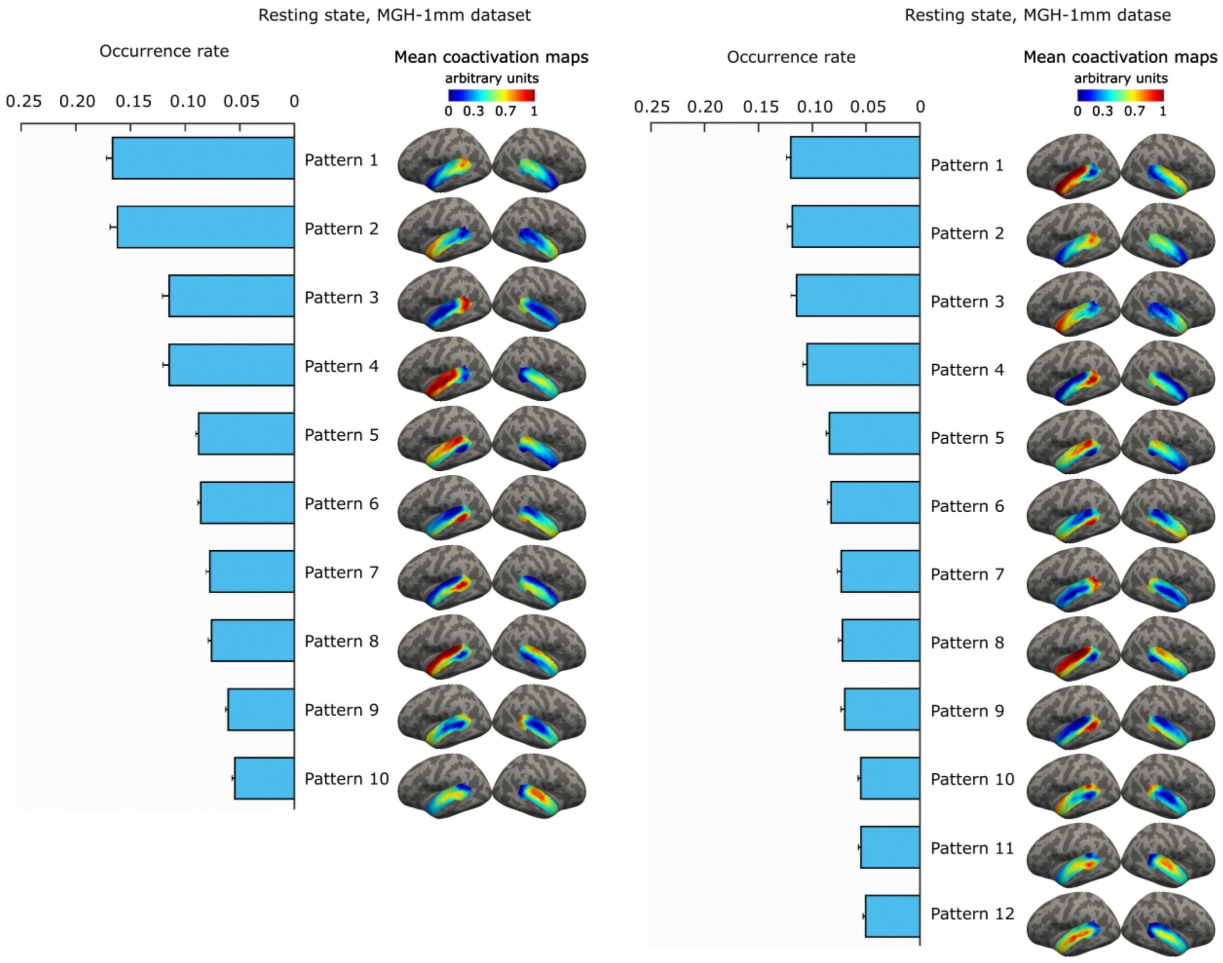


**Figure S3**: Occurrence rates and coactivation maps for 10 and 12 AC pattern solutions derived from the resting-state MGH-1mm data. INSCAPE algorithm was used to generate the group templates of the AC coactivation patterns from 7T fMRI data of 30 healthy participants. The group-level coactivation maps for the patterns were computed as an average over the fMRI time frames assigned to the same cluster across participants. The AC coactivation patterns were ranked by their co-occurrence rates in descending order. Error bars indicate the standard error of the mean (SEM).

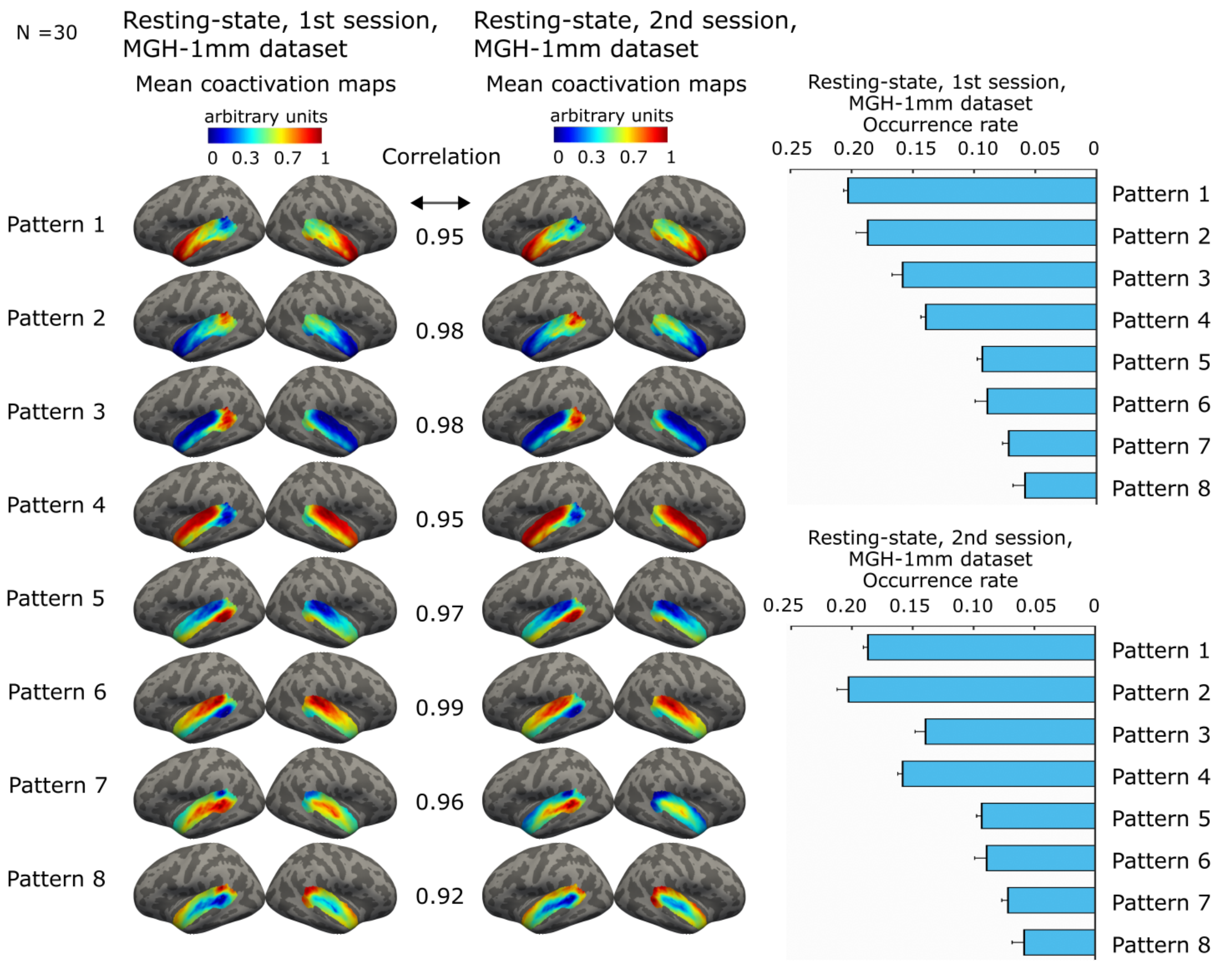


**Figure S4**: Reproducibility of the AC patterns between resting-state fMRI sessions of the MGH-1mm participants. INSCAPE algorithm was used to generate the group templates of the AC coactivation patterns and the corresponding coactivation maps and occurrence rates separately from the first and second resting-state sessions. The coactivation maps were arranged in descending order based on their spatial correlation with the maps derived from the entire resting-state dataset (Fig. 2 in the main text). The correlation values indicate the Spearman correlation between the corresponding patterns of the two datasets. All correlation values were statistically significant ($p<0.001$ for all patterns). The Pearson correlation between the occurrence rates was 0.96 ($p<0.001$). Error bars indicate the standard error of the mean (SEM).

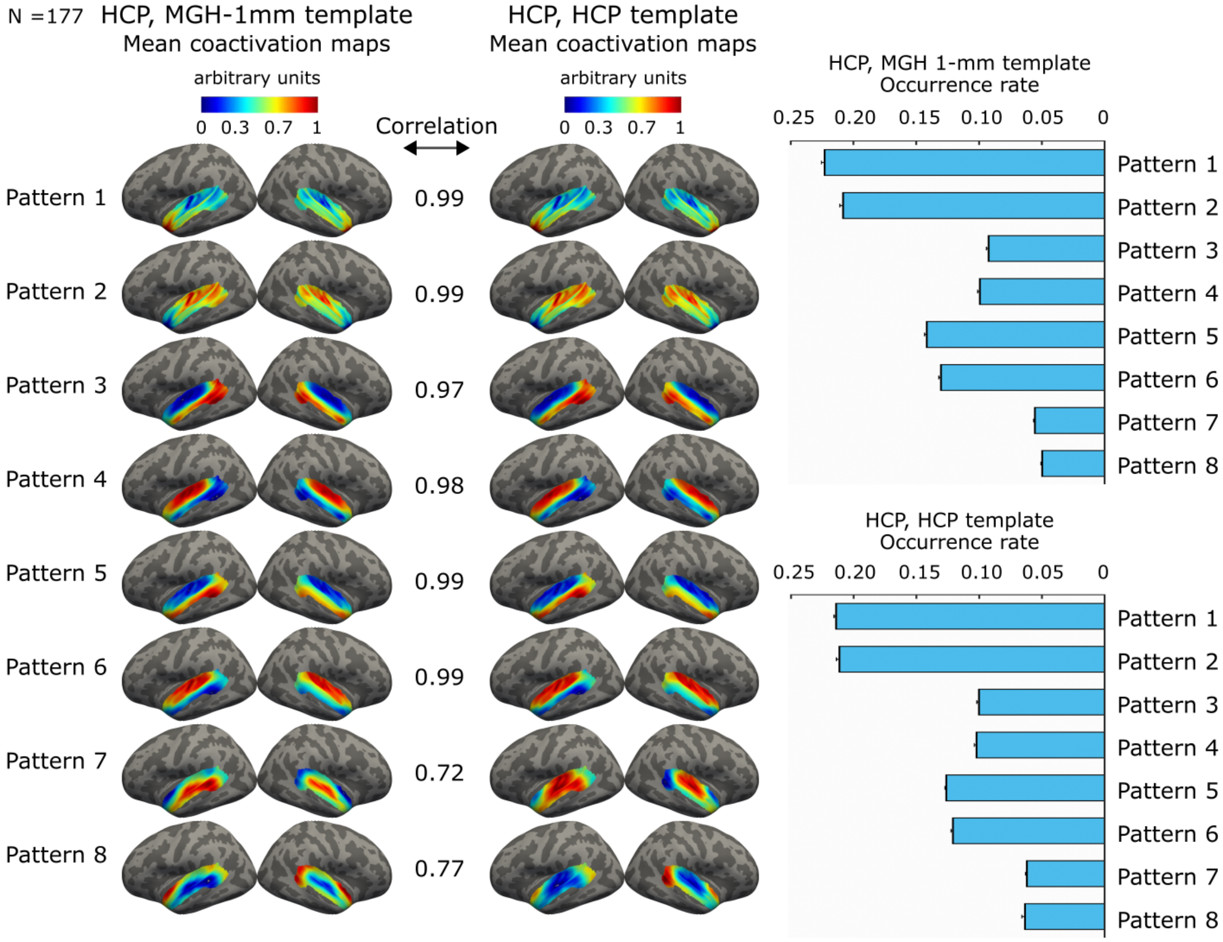


**Figure S5**: Generalizability of the AC template patterns between datasets. The mean coactivation maps and occurrence rates of the AC patterns computed from the HCP resting-state data using the group templates determined from the MGH-1mm data and from the HCP data. The coactivation maps are arranged in descending order based on their spatial correlation with the maps derived from the entire resting-state MGH-1mm dataset (Fig. 2 in the main text). The correlation values indicate Spearman correlation between the corresponding patterns of the two datasets. All correlation values were statistically significant ($p<0.001$ for all patterns). The Pearson correlation between the occurrence rates was 0.99 ($p<0.001$). Error bars indicate standard error of the mean (SEM).

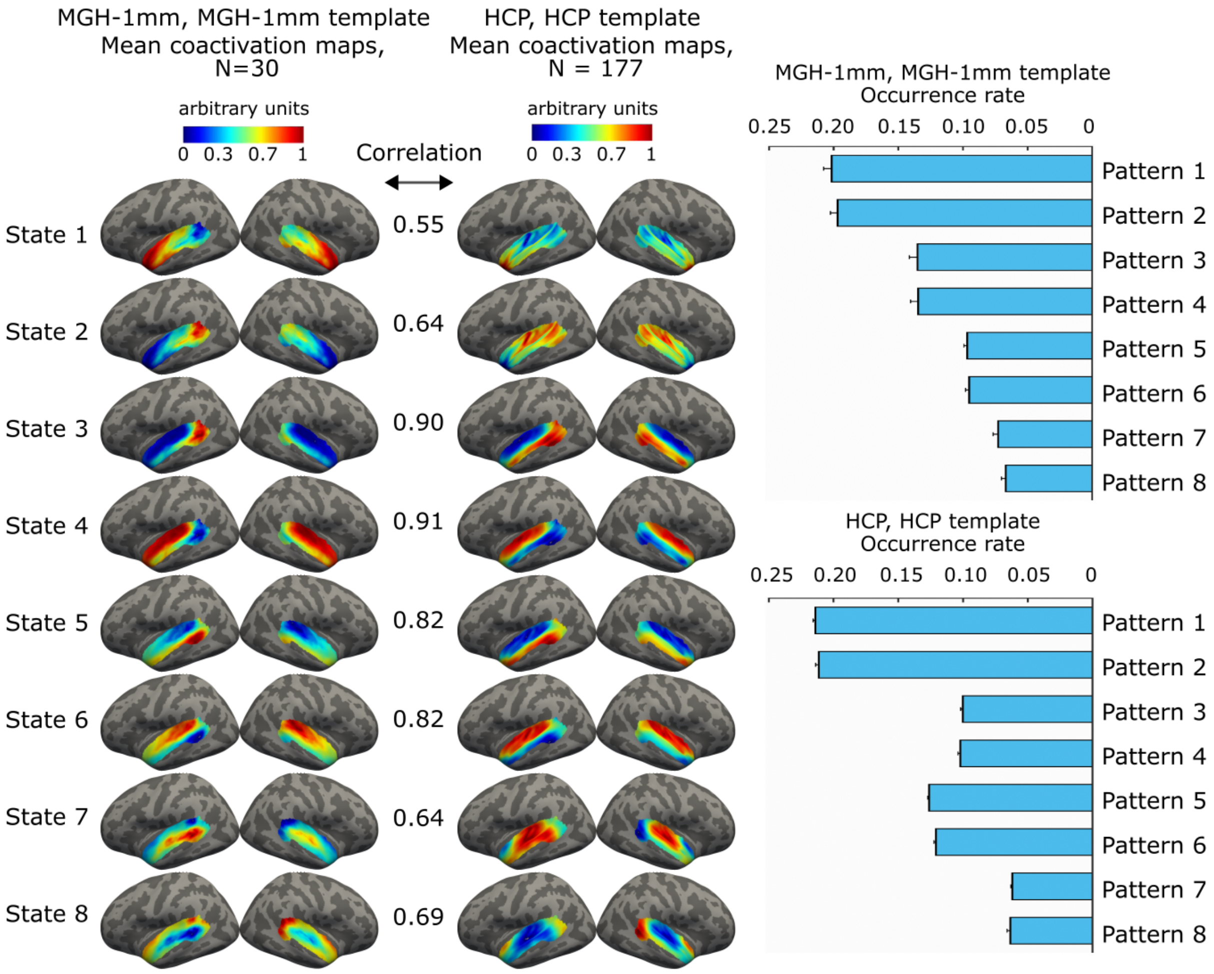


**Figure S6**: Generalizability of the AC pattern coactivation maps between datasets. A) The mean coactivation maps and occurrence rates of the AC patterns computed from the resting-state MGH-1mm data and HCP data. The MGH-1mm maps were determined using the group templates from the same MGH-1mm data. The HCP maps were determined using the group templates from the same HCP data. The coactivation maps are arranged in descending order based on their spatial correlation with the maps derived from the entire resting-state MGH-1mm dataset (Fig. 2 in the main text). The correlation values indicate the Spearman correlation between the corresponding patterns of the two datasets. All correlation values were statistically significant ($p<0.001$ for all patterns). The Pearson correlation between the occurrence rates was 0.91 ($p<0.002$). Error bars indicate the standard error of the mean (SEM).

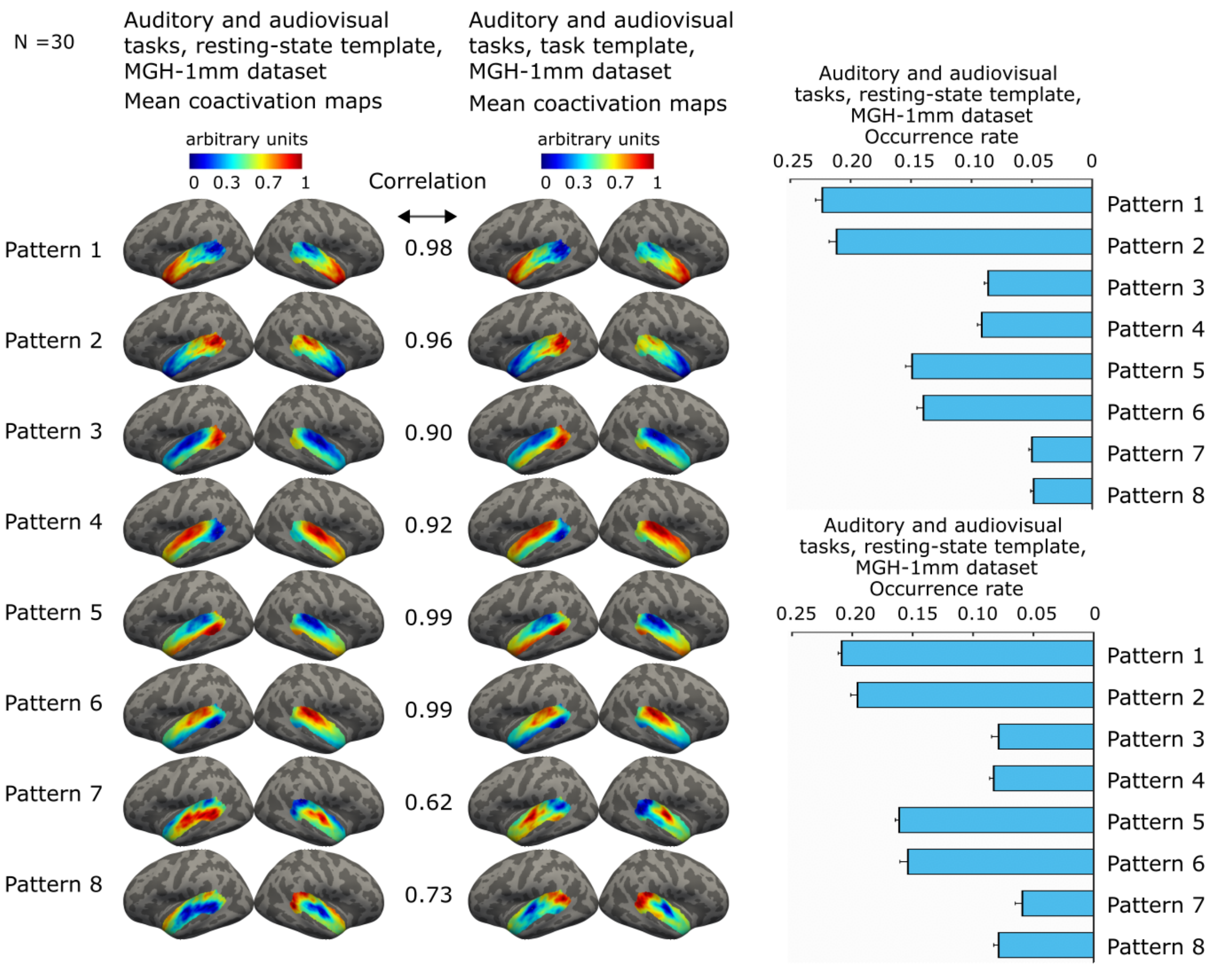


**Figure S7**: Generalizability of the resting-state AC pattern templates to the task data. The mean coactivation maps and occurrence rates of the AC patterns computed from the task fMRI data using the group templates determined from the resting-state data and from the task data. The correlation values indicate Spearman correlation between the corresponding patterns of the two datasets. All correlation values were statistically significant ($p<0.001$ for all patterns). Error bars indicate standard error of the mean (SEM).
